# Size matters - the impact of nucleus size on results from spatial transcriptomics

**DOI:** 10.1101/2022.03.31.486657

**Authors:** Elyas Mohammadi, Katarzyna Chojnowska, Michał Bieńkowski, Anna Kostecka, Magdalena Koczkowska, Michał A. Żmijewski, Marcin Jąkalski, Martin Ingelsson, Natalia Filipowicz, Paweł Olszewski, Hanna Davies, Justyna M. Wierzbicka, Bradley T. Hyman, Jan P. Dumanski, Arkadiusz Piotrowski, Jakub Mieczkowski

## Abstract

Visium Spatial Gene Expression (ST) is a method combining histological spatial information with transcriptomics profiles directly from tissue sections. The use of spatial information has made it possible to discover novel modes of gene expression regulations. However, in the ST experiment, the nucleus size of cells may exceed the thickness of a tissue slice. This may, in turn, negatively affect comprehensive capturing the transcriptomics profile in a single slice, especially for tissues having large differences in the size of nuclei. Here, we defined the effect of Consecutive Slices Data Integration (CSDI) on results from spatial transcriptomic study of human postmortem brains. CSDI can be applied to investigate consecutive sections studied with ST in the human cerebral cortex, avoiding misinterpretation of spot clustering and annotation, increasing accuracy of cell recognition as well as improvement in uncovering the layers of grey matter in the human brain.

## Introduction

The spatial transcriptomics concept has been introduced as a combination of massively parallel sequencing and microscopic imaging^1^. This method is an attractive approach in studies of normal development and in clinical translational research. Visium Spatial Gene Expression (ST) is one of the technologies developed around this concept. ST is a next-generation molecular profiling method dedicated to unraveling the transcriptomic architecture of the tissue. The application of ST for mapping the transcriptome with the morphological context has been proven successful in many fields^2^.

Although ST is a powerful new technique for capturing patterns of spatial distribution of gene expression, it also has a drawback of its design. A Visium Gene Expression slide consists of two or four tissue-capture areas (6.5 mm x 6.5 mm), divided into 4992 spots, each 55 μm in diameter. Every spot contains oligonucleotide probes with unique sequence barcodes that encode spatial information in gene expression data^2^. Due to their size, spots may encompass the expression profiles of several cells. Consequently, this diminishes the accuracy of distinguishing neighboring cell types. This can be addressed by several methods^3,4^, including integration with other single-cell analyses^5^. The most popular methods for the integration rely on so called anchors, which represent similar gene expression patterns.

The importance of the anchor-based data integration in distinct single-cell modalities (i.e., spatial transcriptomics and single nucleus RNA sequencing data [snRNA-seq]) has been investigated previously^5^. However, the application of Consecutive Slices Data Integration (CSDI) in ST analysis using the anchor-based approach remains unexplored. We investigated the effects of CSDI on spot clustering and cell-type annotation using both snRNA-seq and ST technologies in human cerebral cortex samples. By applying the CSDI to ST, we aimed to evaluate whether a single slice of tissue would be sufficient for ST analysis or whether consecutive slices would be required. We found that without CSDI, the pattern of obtained spot clusters between consecutive slices is inconsistent, and the cell-type annotation does not match the microscopic characterisation of the slice. These issues were resolved by employing CSDI, and layer-structure of grey matter of the human brain was unveiled.

## Results

We conducted spatial gene expression analysis in human postmortem, fresh frozen tissue sections. Two anatomical regions, the Orbitofrontal Neocortex (ON) and the Temporal Neocortex (TN) from two adult male donors were investigated (Figure 1). From each region of both subjects, one pair of consecutive slices (eight slices in total) were prepared. We cut the ON and TN tissue into 10-12 μm thick sections. Each sample was sequenced to a median depth of 187 million reads, corresponding to a mean of 3300 unique molecular identifiers (UMIs) and a mean of 2058 genes per spot.

**Figure 1.**
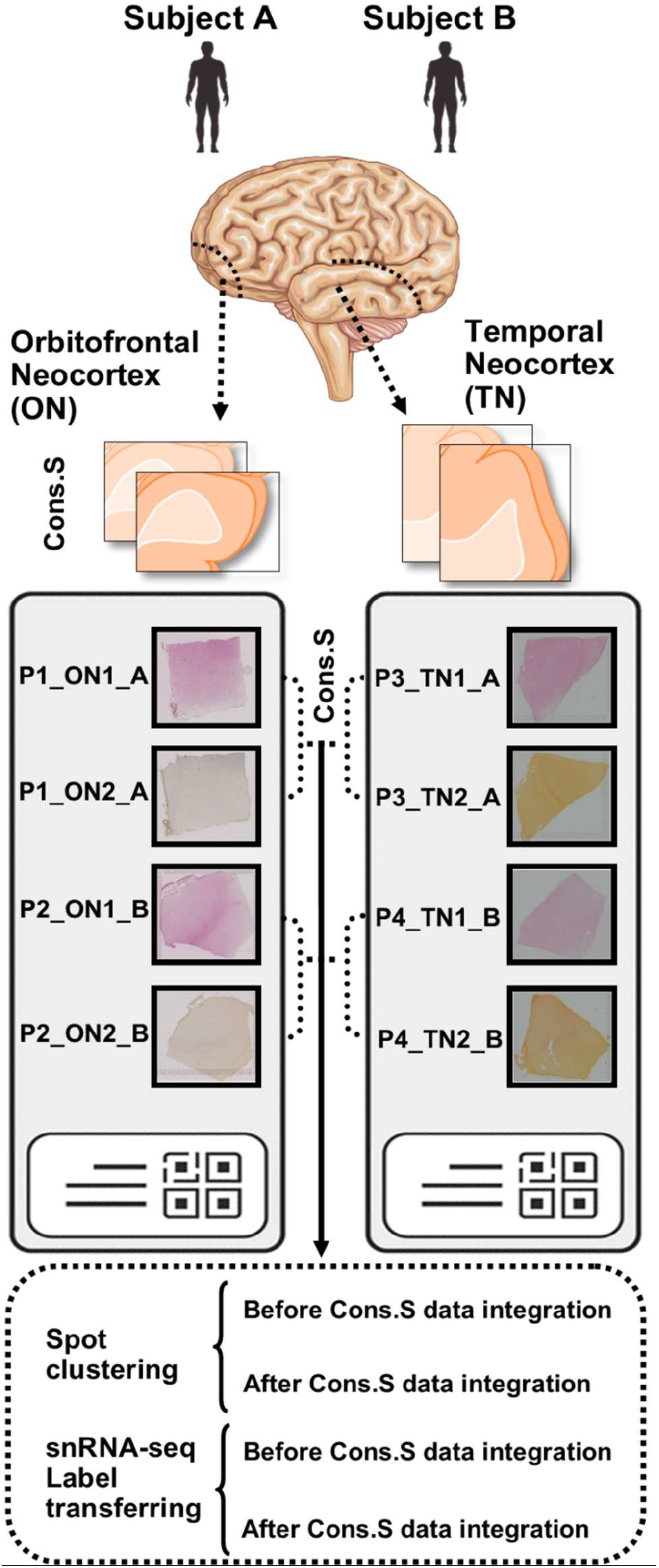
Schematic view of experimental workflow and *in silico* analytical pipeline. Four pairs (P1–P4) of consecutive slices of human postmortem brains were obtained from two distinct anatomical locations (ON and TN). Two types of computational analyses were performed (spot clustering and snRNA-seq label transferring) and further divided into additional subtypes (before and after CSDI for each category). The names of slices are as follows: P, pairs of consecutive-sections; ON (1–2), Orbitofrontal Neocortex (section number in pair of consecutive slices), TN (1–2): Temporal Neocortex (section number in pair of consecutive slices); A and B: two studied subjects; Cons. S: consecutive sections.

### Identifying distinct cell types and their annotation using single tissue section

Figure 2A shows the distinction between the grey matter (GM) and the white matter (WM). The border between GM and WM was established histologically based on cellular composition and arrangement (Figure 2A, Z1, Z2, and Z3). We used an unsupervised method to investigate whether categorizing the ST spots based on their transcriptomics profile could represent structural layers of the brain. Subsequently, we compared the obtained groups with histological images of tissue slices to assess the obtained clusterization and classification (Figure 2A). Thus, we confirmed the general consistency of GM and WM patterns revealed by histologic and transcriptomic methods.

**Figure 2.**
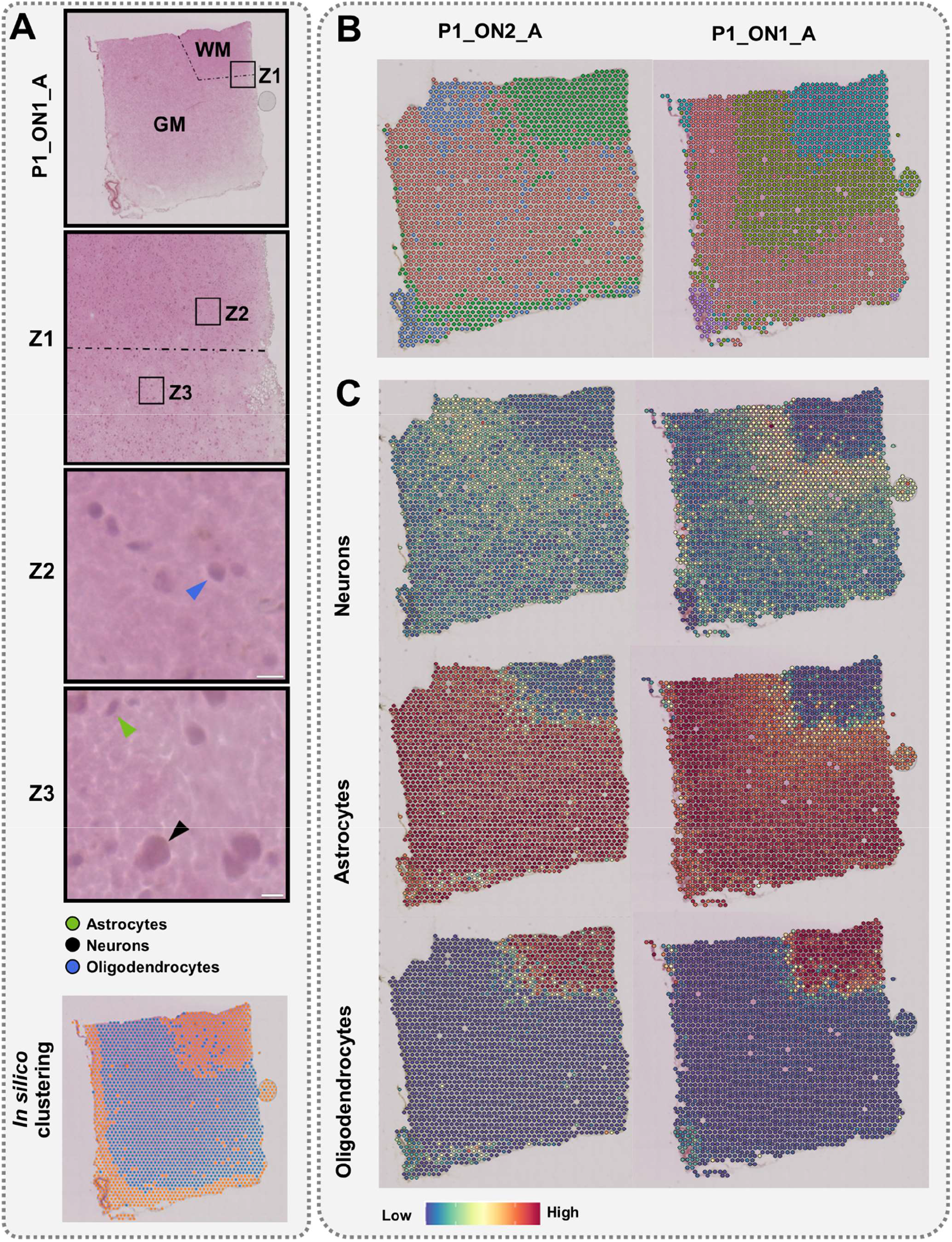
Results from spatial transcriptomics analysis using single tissue sections. **Panel A**; Top, histological image of orbitofrontal neocortex (ON) with marked white matter (WM) and grey matter (GM); **Z1,** zoomed-in image of the border between WM and GM; **Z2,** Blue arrow points to an oligodendrocyte nucleus; **Z3,** Black and Green arrows represent nuclei of neurons and astrocytes, respectively. In Z2 and Z3, white scale bars represent 10 μm. Bottom, unsupervised classification of spots. **Panel B;** The ST spots clustering before CSDI. **Panel C;** Label transferring before CSDI. The name of the sample encodes number of the section (P1/P2), number of slice (ON1/ON2), and patient id (A/B).

We performed spot clustering using the steps recommended by Satija et al. 2015^6^ in order to cluster the spots with similar expression profiles, and distinguish distinct cellular layers. The resulting clusters revealed the separation of subcortical WM and cortical GM. More detailed morphological layers of the brain were also unveiled through the more detailed clustering (Figure 2B). Considering the expected similarity of architecture between two consecutive slices of the cerebral cortex, we should observe the very similar pattern of clusters. However, this consistency was vague, and the layered structure of GM in P1_ON2_A could not be observed (Figure 2B). We observed that although, the use of a single section of brain tissue with the ST method can be informative, it may also have critical limitations in spot clustering. To overcome this, we decided to use external gene expression data set and anchor-based integration method.

To better understand the identified brain layers, we integrated the measured expression profiles with a previously described snRNA-seq dataset^7^. As single nucleus profiles contain greater number of genes than in our ST profiles, the integration of these two data sets allowed us to perform the spot annotation more precisely. Using predefined cell-type annotations in snRNA-seq—including oligodendrocytes, astrocytes, and neurons—the ST spots were labeled (see “Materials and methods” for details). The pattern of transferred labels is shown in P1_ON1_A and P1_ON2_A as an example (Figure 2C). The locations of oligodendrocytes and astrocytes were primarily identified in WM and GM, respectively, in line with brain structure (Figure 2A). However, we could not confidently annotate neurons in GM, which is incompatible with histology (Figure 2A). In summary, a single slice of brain tissue using the ST method is informative but has limitations in distinguishing cell types using label transferring as well as in spot clustering.

### The effects of CSDI on identifying distinct cell types and their annotations

Stuart et al. 2019^5^ developed the CSDI to correct the transcriptomic profiles of consecutive slices using anchors representing spots with similar gene expression profile from two consecutive slices. This is used to pair spots from the two slices. At the same time, the transcriptomic differences between pairs of spots in anchors are used to correct datasets from both consecutive sections.

We decided to apply the CSDI method due to the heterogeneity of the brain in terms of size of nuclei among different cell types (Figure 3). On average, the size of a nucleus from a neuron in GM (about 20 μm) is much larger than the thickness of tissue section (10-12 μm). Consequently, a single tissue section will encompass only part of nuclei for essentially all neurons present in the studied sample. This restriction will also apply to other smaller nuclei, although to a lesser extent. Thus, for all cells present in a studied brain tissue, it will cause partial loss of transcriptomic signals. Taking “P1_ON1_A” and “P1_ON2_A” as consecutive slices of ON as an example, we could identify all the morphological layers of the brain^8^ only after CSDI (Figure 4A and Supplementary Figure 1). In conclusion, CSDI can resolve the issue of inconsistency of the pattern of clustering between consecutive slices. Moreover, by applying the same parameters (see “Materials and methods” for details), we can identify more neuronal layers in GM^9^ (Figure 4A and Supplementary Figure 1).

**Figure 3.**
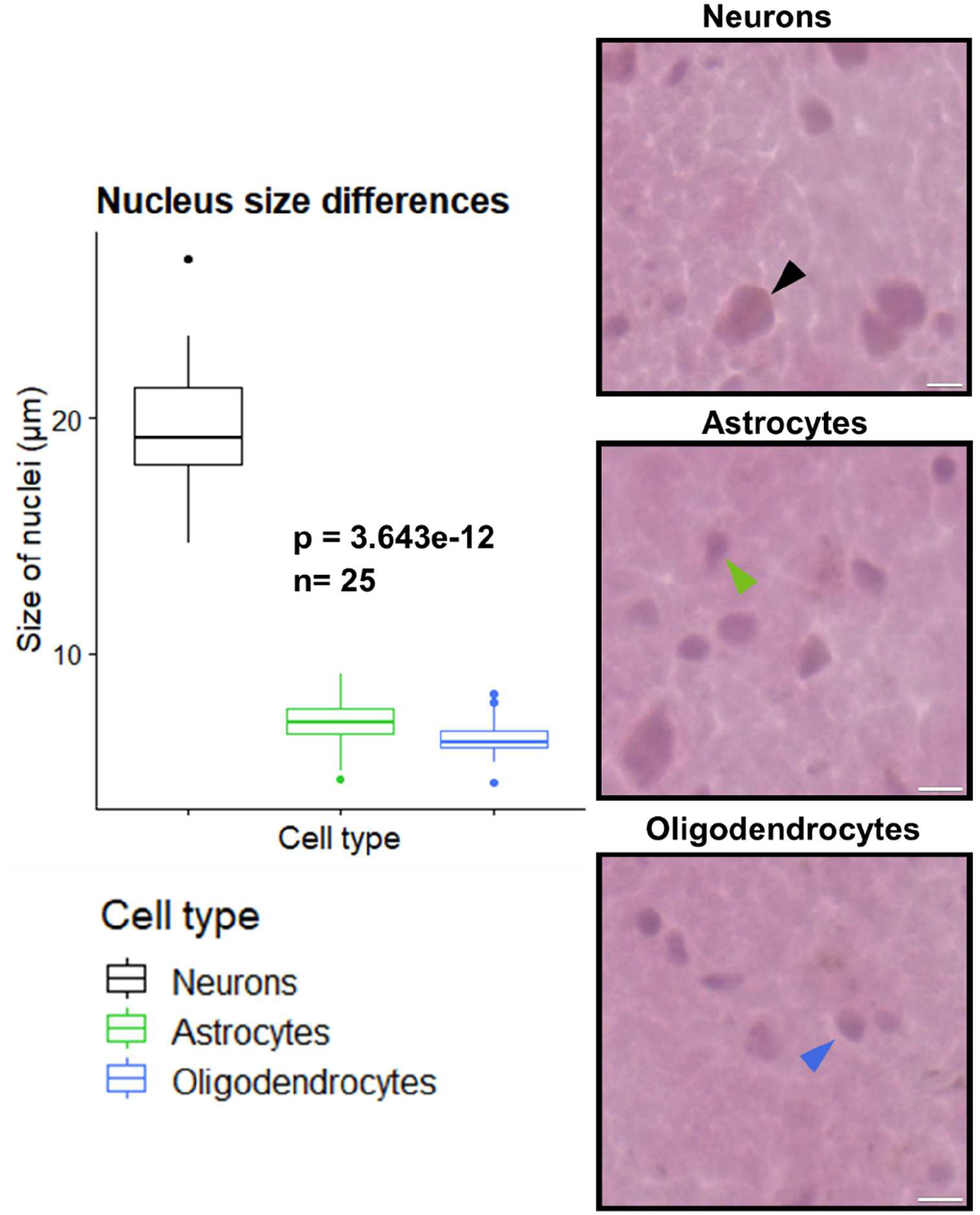
The nucleus size heterogeneity among investigated cell types. Left: The boxplot shows significant differences in size of nuclei in the human cerebral cortex (Kruskal-Wallis test). Three panels on the right; Arrows represent examples of nuclei for three cell types in the P1_ON1_A sample. Scale bars denote 10 μm.

**Figure 4.**
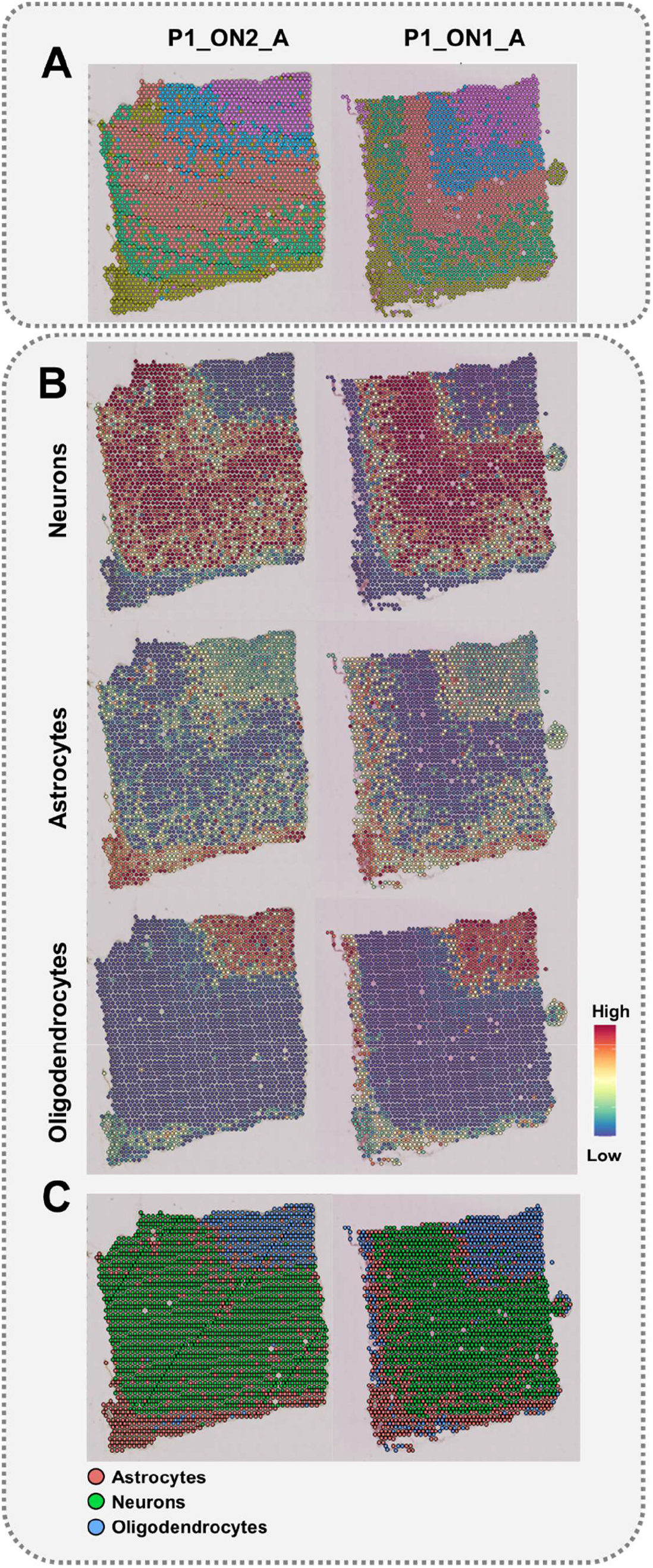
The CSDI method can improve the clustering and spot annotation of consecutive slices. **A)** Clustering after CSDI unveiled the GM layers and resolved the inconsistency between the pattern of clusters in consecutive slices. **B)** snRNA-seq Label transferring after CSDI. The annotation probability is shown as a scheme for three different cell types. **C)** The classification of spots through label transferring is shown by considering the highest probability of annotation for each spot after CSDI.

Cell bodies of neurons are mainly found in the GM (Figure 2A). However, during our label transferring, the spots marked as neurons received weak probability values in the GM (Figure 2C). This is an important concern, which led us to hypothesize that using information from a single section of tissue may lead to inaccurate interpretation of clusters and cell types. Our approach to transferring cell labels from snRNA-seq to ST before and after CSDI revealed different results, which provides support for the above hypothesis. These differences are much more pronounced in GM, where the spots recognized as neurons, or astrocytes are the dominating cell types (Figure 4B). Accordingly, we compared the annotations with consideration for the size of nuclei and the structural layers of the brain (WM and GM). Prior to CSDI, the annotation of neurons and astrocytes received low and high probabilities, respectively, in the area of the GM, while the nuclei of oligodendrocytes were mostly visible in the WM (Figure 2C and Supplementary Figure 2). Interestingly, after CSDI, the likelihood of the annotation of neurons was increased due to the gain of neuronal transcriptomic profiles (Figure 4B), which is consistent with the histological imaging (Figure 2A). It is noteworthy that we did not observe any changes in the probability of annotation for oligodendrocytes in the WM before and after CSDI, which is also in agreement with histological structure of the cerebral cortex. Among the transferred labels from snRNA-seq to ST, neurons and astrocytes are mainly available in GM. Consequently, no signal fluctuation will occur before and after CSDI in WM. In both situations, the WM would preferentially be annotated with oligodendrocytes (Figure 2C, Figure 4B, and Supplementary Figure 2).

The differences between annotations obtained for the spots before and after CSDI can be attributed to the fact that the size of neuronal nuclei is much bigger than astrocytic nuclei^10^ (Figure 3). Their size is actually larger than the thicknesses of the tissue sections used in the ST protocol. Accordingly, a single slice will capture incomplete transcriptomic neural context. CSDI provides a robust means of rectification of this misinterpretation. Hence, the corrected signals of all types of nuclei can be obtained. Consequently, the label transferring from snRNA-seq to ST is made consistent with the histological findings (Figure 2A). Ultimately, one can study the spatial distribution of different cell types more precisely.

We evaluated the results from two independent spot-categorization methods used in this study: label transferring and spot clustering. Hence, we classified the spots using the transferred labels (Figure 4C and Supplementary Figure 3) and compared them with the spot clustering results represented in Figure 4A. We observed that the green cluster in Figure 4C represents the three distinguished neural layers in Figure 4A (blue, red, and green clusters). Similarly, the blue cluster in Figure 4C represents oligodendrocytes in Figure 4A (purple cluster). Through determining the locations of neurons and oligodendrocytes in both methods, we demonstrated that the results from both spot-categorization methods are consistent with the histological images (Figure 2A).

## Discussion

We studied the impact of the CSDI method on spatial gene expression analysis and evaluated the effect of CSDI on the improvement of clustering and label transferring^5^. The application of CSDI was motivated by the two issues we observed in the results of the basic spatial transcriptomic analysis. Firstly, in the GM, we observed inconsistencies between the patterns of clusters in consecutive slices (Figure 4A). Secondly, we failed to recreate the expected layered structure of GM (Figure 4A). According to the study conducted by Maynard et al. 2021^9^, data correction in consecutive slices was performed using the data-refinement step of Space Ranger. Hence, the spatial topography of gene expression in the human dorsolateral prefrontal cortex was defined. The pattern of determining clusters was consistent in all pairs of consecutive slices, a phenomenon we observed only after applying the CSDI. Moreover, using CSDI, we distinguished cortical and subcortical WM layers. Thus, we showed that the expected consistency of the pattern of clustering between consecutive slices can be achieved with CSDI similar to Space Ranger.

We investigated the results of the clustering and label transferring, with and without CSDI utilization. Simultaneously, we compared the consistency of the results obtained with the topographic organization of the cerebral cortex. We observed the improvement of clustering and the label transferring after applying CSDI. The superior performance of using CSDI is likely related to the size of nuclei in different cell types as the determining parameter. The sizes of the nuclei of certain neurons are much larger than the nuclei of astrocytes and oligodendrocytes (e.g., neurons from the pyramidal layer (Figure 3) of the cerebral cortex)^10^. In the human brain, the size of neural nuclei may exceed the tissue thickness recommended in the cryosectioning step of the ST protocol (10 μm)^11^. This may jeopardize capturing the whole transcriptomics profile using a single slice.

Experiments involving tissue sections or entire organ cross-sections from small animals are virtually free from the risk of losing the transcriptomics content of cells. For example, in mice, the average diameter of neural soma derived from the cortical pyramidal layer does not exceed 10 μm^12^. Hence, our approach is specifically applicable to tissues composed of cells with nuclei sizes exceeding the minimum thickness of the section required for the spatial transcriptomics experiment.

We used a combination of ST and snRNA-seq technologies to unveil the cerebral-cortex structure and related cell types. The ST preserves the spatial location of gene expression. However, its resolution at the level of the spot, as well as in terms of the number of captured genes, is nominally lower than the single-nuclei/single-cell transcriptomics^13^. The lower resolution results from the size of spots in ST expression slides (55 μm in diameter). Accordingly, each spot may encompass the transcriptomic profiles of multiple cells. The ST data integration with snRNA-seq/scRNA-seq is considered a deconvolution method to unravel the underlying cell types in each ST spot. In this context, using snRNA-seq has advantages over scRNA-seq. This is because the process of tissue cryopreservation ruptures the cell membranes; however, nuclear membranes remain intact during the freeze-thaw cycle^14^. Furthermore, it has been shown that the RNA-seq of single nuclei is viable and highly representative of transcriptional profiles from the entire cells. This fact is specifically relevant to postmortem brain samples after long-term storage at −80 °C^14^. Additionally, using the protease to dissociate cells in scRNA-seq leads to the activation of the crucial, immediate early genes that may skew experimental data^15^. Hence, we utilized the prelabeled snRNA-seq to deconvolute the ST spots.

To confirm the deconvolution of ST spots and defined cell types, we compared our annotation with neuropathological findings. Astrocytes play a vital role in delivering energy to neurons via the astrocyte-neuron lactate shuttle^16^. Hence, astrocytic nuclei are spatially located beside perikarya (Figure 2A), mostly placed in the GM^17^. In Figure 4B, the GM is annotated for both neurons and astrocytes, corresponding to the previous findings^16^. According to the shape of oligodendrocytic nuclei—which are round with visible halos^18^—the annotation of WM for oligodendrocytes corresponds with the expected normal morphology of the brain cross-section^19^ (Figure 3). These concepts are consistent with our histological (WM and GM) (Figure 3) and cell-type (astrocytes, neurons, and oligodendrocytes) (Figure 4C) classifications.

An alternative solution to resolve the low resolution of the ST method is to decrease the size of barcoded spots in gene expression slide glasses. However, as we addressed in our study, ST results would be affected by the size of neural nuclei because the origin of the problem is not the sizes of spots but the thickness of the tissue slices. Accordingly, by decreasing the sizes of capture spots, deconvolution methods may no longer be required anymore; however, the need for CSDI remains.

In summary, the transcriptomic profiles of ST consecutive slices may need to be corrected prior to further analysis. Correcting the datasets simply for the depth of sequencing using normalization methods (e.g., log normalization) cannot remove all the unknown batch effects of consecutive slices. Data correction can be performed during the data-processing step by Space Ranger using the *spaceranger aggr* function or during the analysis steps using CSDI. In Space Ranger, the transcriptomic profile of consecutive slices will be aggregated, normalized to the same sequencing depth. Then, the feature-barcode matrices and the analysis of the combined data can be recomputed. In CSDI, the spots with similar transcriptomics profiles in two datasets will be anchored. Using the anchors, the transcriptomics profile of consecutive slices will be corrected, and one can proceed with the downstream analysis. Consequently, the results of clustering and annotation will be improved after data correction. Therefore, more trustable biological findings can be achieved.

## Materials and methods

### Data acquisition

We utilized the modified 10x Genomics Visium Spatial Gene Expression method to analyze the profiles of consecutive sections from fresh-frozen brain tissues. Accordingly, we used the orbitofrontal (ON) and temporal neocortex (TN) samples from two subjects. Tissue specimens were provided by Harvard University and Massachusetts Alzheimer’s Disease Research Center and all experimental procedures were conducted in accordance with Independent Bioethics Committee for Scientific Research at Medical University of Gdansk (consent No. NKBBN/564-108/2022). The brain-tissue slices were placed onto a Visium Gene Expression slide (10x Genomics) and fixed according to the 10X Genomics protocol (doc. CG000239 Rev. C). Next, the slides were divided into two via a piece of silicone gasket. Subsequently, we stained the tissue by two methods—hematoxylin and eosin, and hematoxylin and Congo red—to detect eventual amyloid deposits. We imaged the slides at 20x magnification using brightfield settings (Olympus cellSens Dimension software). Afterward, the tissue was permeabilized, using conditions according to manufacturer protocol. The mRNA was released and bound to spatially barcoded capture probes on the slide. Next, cDNA was synthesized from captured mRNA, and sequencing libraries were prepared. Samples were loaded and pooled according to the protocol (doc. CG000239 Rev C) and sequenced in the standard Illumina pair-end constructs, using Illumina’s NextSeq 550 System.

### Visium data processing

Four pairs (consecutive slices) of ST raw data (BCL files) from two postmortem brain samples were converted to fastq files using 10x Genomics software Space Ranger version 1.2.1 and its *spaceranger mkfastq* function. Subsequently, reads were aligned to the human genome-reference sequence (GRCH38) using the STAR method, and spatial feature counts were generated using the *spaceranger count* function. Because an inverted microscope was used for imaging, all images were flipped horizontally prior to being applied to the Space Ranger.

### Data preprocessing and normalization

All outputs from *spaceranger count* were read as Seurat objects using *Load10X_Spatial* function of Seurat version 4.0.3. Prior to data normalization, the percentages of mitochondrial genes were calculated by the *PercentageFeatureSet* function. Then, the spots with a number of spatial features of more than 7000 and less than 200 were removed; spots that encompassed more than 15% of mitochondrial genes also were omitted from the downstream analyses. Standard normalization was performed using the *NormalizeData* function and the *LogNormalize* method using default parameters. Variable features for each object were determined using the *FindVariableFeatures* function and *VST* method. Next, the data were scaled and regressed out for the percentage of mitochondrial genes using the *ScaleData* function.

### Dimensionality reduction and clustering

Dimensionality reduction was completed using the *RunPCA* function. Prior to clustering, nearest neighbors were determined by the *FindNeighbors* function with default parameters. After this, the *FindClusters* function determined the clusters by a shared nearest-neighbor (SNN) modularity optimization-based clustering algorithm (The resolution was arbitrarily set to 0.3).

### Consecutive slices data integration

Dimensionality reduction for consecutive slices was completed jointly through diagonalized canonical correlation analysis (CCA). Using the *FindIntegrationAnchors* function, Mutual Nearest Neighbors (MNNs) were found in this shared low-dimensional space and were termed anchors (for more details, see Stuart et al., 2019^5^). The *IntegrateData* function was considered for CSDI using precomputed anchor sets. The integrated consecutive slices were saved as transcriptomics-corrected objects. The same workflow for dimensionality reduction and clustering was applied to integrated objects. Finally, the eight Seurat objects (four pairs of consecutive slices) before and after CSDI (16 in total) were saved as RDS files to be compared from different perspectives.

### Label transferring from snRNA-seq to ST

The previously annotated snRNA-seq dataset was obtained from scREAD, a publicly available snRNA-seq database^20^ (https://bmbls.bmi.osumc.edu/scread/). The causes of death for the two donors were Alzheimer’s disease (Subject A) and stroke (Subject B). Hence, snRNA-seq profiles with AD01104 and AD01102 scREAD data IDs for Alzheimer’s and non-Alzheimer’s disease were retrieved^7^. The raw-sequencing data and the digital-expression matrices obtained using the 10x Genomics software Cell Ranger are available in the NCBI’s Gene Expression Omnibus (GSE129308) and are accessible through the GEO Series accession number GSM3704357-GSM3704375^21^.

Data normalization and dimensionality reduction with the same parameters as the ST data were conducted for the two snRNA-seq datasets. By considering snRNA-seq as our reference and ST data as query datasets, anchors were found, and precomputed cell labels were transferred using the *FindTransferAnchors* (identifying shared cell/spot states present across different datasets) and *TransferData* functions, respectively. Label transferring and clustering were completed twice for each of the ST objects—once before and once after CSDI—to investigate the effect of CSDI on label transferring and clustering.

## Data availability

The snRNA-seq data used in this study are publicly available at the scREAD database (https://bmbls.bmi.osumc.edu/scread/) under AD01104 and AD01102 scREAD data IDs. Additionally, the snRNA-seq raw data and the digital-expression matrices obtained using the 10x Genomics software Cell Ranger are available in the NCBI’s Gene Expression Omnibus (GSE129308) and are accessible through the GEO series accession numbers GSM3704357-GSM3704375. The spatial transcriptomics data used in this study are privately available at the GEO data repository under the GSE184510 accession number. These data can be available from the authors upon a reasonable request.

## Code availability

The scripts used in this study are developed by the R programming language version 4.1.0 and have been deposited in a public Github repository (https://github.com/ElyasMo/ST_snRNA-seq).

## Acknowledgments

This work was supported by the Foundation for Polish Science under the International Research Agendas Program financed from the Smart Growth Operational Program 2014–2020 (Grant Agreement No. MAB/2018/6). This study was also partly supported by grants from the Swedish Cancer Society, the Swedish Research Council, Hjärnfonden, and Alzheimerfonden, to JPD and P30AG062421/AG/NIA NIH HHS/United States, to BTH.

## Author contributions

**Conceptualization:** AP, JPD, JM, KC, EM, MB, PO

**In silico data analysis:** EM, AP

**Formal analysis:** EM, KC, AP

**Funding acquisition:** JPD, AP

**Investigation:** KC, EM, JM, AK, MK

**Project administration:** JPD, AP, NF, HD

**Resources:** JPD, AP, BH, MI, MZ, JW,

**Software:** EM, JM, MZ

**Supervision:** JM, AP, JDM, MB, MJ

**Validation:** KC, EM, MB, AK, MK

**Writing-original draft:** EM, KC, JM, AP

**Writing-review & editing:** EM, KC, JM, JPD, MB, AK, BH, MI, AK, MK, MJ, NF, MZ, JW, PO, HD

## Competing interests

All authors declare no competing interests.

## Supplementary data

**Supplementary Figure S1.** Improvement of the spot clustering through CSDI. **Panel A:** the expected consistency of the pattern of spot clusters between consecutive slices is missed. **Panel B:** the consistency of clustering after consecutive slices data integration is improved.

**Supplementary Figure S2.** Improvement of the label transferring by CSDI. The annotation probability is shown as a scheme for three different cell types. Label transferring from snRNA-seq to ST consecutive slices is shown before and after data integration. **Before Integration:** the probability of spot annotations for neurons is not compatible with tissue histology. **After integration:** the probability of the presence of neurons increased in the GM of the cerebral cortex.

**Supplementary Figure S3.** A comparison between the pattern of spot classification after CSDI through spot clustering and label transferring methods. The consistency between the results from these two methods, which is compatible with the histological information of tissue slices, supports the accuracy of the spot categorization.

## Notes

https://bmbls.bmi.osumc.edu/scread/

https://github.com/ElyasMo/ST_snRNA-seq

### Competing Interest Statement

The authors have declared no competing interest.

## References

1 Rao, A., Barkley, D., França, G. S. & Yanai, I. Exploring tissue architecture using spatial transcriptomics. Nature 596, 211–220, doi:10.1038/s41586-021-03634-9 (2021).

2 Asp, M., Bergenstråhle, J. & Lundeberg, J. Spatially Resolved Transcriptomes—Next Generation Tools for Tissue Exploration. BioEssays 42, 1900221, doi:https://doi.org/10.1002/bies.201900221 (2020).

3 Dong, R. & Yuan, G.-C. SpatialDWLS: accurate deconvolution of spatial transcriptomic data. Genome Biology 22, 145, doi:10.1186/s13059-021-02362-7 (2021).

4 Elosua-Bayes, M., Nieto, P., Mereu, E., Gut, I. & Heyn, H. SPOTlight: seeded NMF regression to deconvolute spatial transcriptomics spots with single-cell transcriptomes. Nucleic acids research 49, e50–e50 (2021).

5 Stuart, T. et al. Comprehensive integration of single-cell data. Cell 177, 1888–1902. e1821 (2019).

6 Satija, R., Farrell, J. A., Gennert, D., Schier, A. F. & Regev, A. Spatial reconstruction of single-cell gene expression data. Nature biotechnology 33, 495–502 (2015).

7 Zhang, Y. et al. Depletion of NK Cells Improves Cognitive Function in the Alzheimer Disease Mouse Model. The Journal of Immunology 205, 502–510 (2020).

8 Zhao, E. et al. Spatial transcriptomics at subspot resolution with BayesSpace. Nature Biotechnology, doi:10.1038/s41587-021-00935-2 (2021).

9 Maynard, K. R. et al. Transcriptome-scale spatial gene expression in the human dorsolateral prefrontal cortex. Nature Neuroscience 24, 425–436, doi:10.1038/s41593-020-00787-0 (2021).

10 von Economo, C. & Triarhou, L. C. Cellular Structure of the Human Cerebral Cortex. (Karger, 2009).

11 10xGenomics. Visium Spatial Gene Expression Optimized Tissues. https://support.10xgenomics.com/spatial-gene-expression/tissue-optimization/doc/specfications-visium-spatial-gene-expression-optimized-tissues (2020).

12 Kim, E. J., Juavinett, A. L., Kyubwa, E. M., Jacobs, M. W. & Callaway, E. M. Three Types of Cortical Layer 5 Neurons That Differ in Brain-wide Connectivity and Function. Neuron 88, 1253–1267, doi:10.1016/j.neuron.2015.11.002 (2015).

13 Navarro, J. F. et al. Spatial Transcriptomics Reveals Genes Associated with Dysregulated Mitochondrial Functions and Stress Signaling in Alzheimer Disease. Iscience 23, 101556 (2020).

14 Grindberg, R. V. et al. RNA-sequencing from single nuclei. Proceedings of the National Academy of Sciences 110, 19802–19807 (2013).

15 Lacar, B. et al. Nuclear RNA-seq of single neurons reveals molecular signatures of activation. Nature communications 7, 1–13 (2016).

16 Siracusa, R., Fusco, R. & Cuzzocrea, S. Astrocytes: Role and Functions in Brain Pathologies. Frontiers in Pharmacology 10, doi: 10.3389/fphar.2019.01114 (2019).

17 Duchatel, R. J., Shannon Weickert, C. & Tooney, P. A. White matter neuron biology and neuropathology in schizophrenia. npj Schizophrenia 5, 10, doi: 10.1038/s41537-019-0078-8 (2019).

18 El Sharouny, S., Shaaban, M., Elsayed, R., Tahef, A. & Abd ElWahed, M. N-acetylcysteine protects against cuprizone-induced demyelination: histological and immunohistochemical study. Folia Morphologica (2021).

19 Hofmann, K. et al. Astrocytes and oligodendrocytes in grey and white matter regions of the brain metabolize fatty acids. Scientific Reports 7, 10779, doi:10.1038/s41598-017-11103-5 (2017).

20 Jiang, J., Wang, C., Qi, R., Fu, H. & Ma, Q. scREAD: A Single-Cell RNA-Seq Database for Alzheimer’s Disease. iScience 23, 101769, doi:https://doi.org/10.1016/j.isci.2020.101769 (2020).

21 Otero-Garcia, M. et al. Single-soma transcriptomics of tangle-bearing neurons in Alzheimer’s disease reveals the signatures of tau-associated synaptic dysfunction. BioRxiv (2020).

