## Supplementary figures and images for "Size matters - the impact of nucleus size on results from spatial transcriptomics"

### Supplementary Figure 1

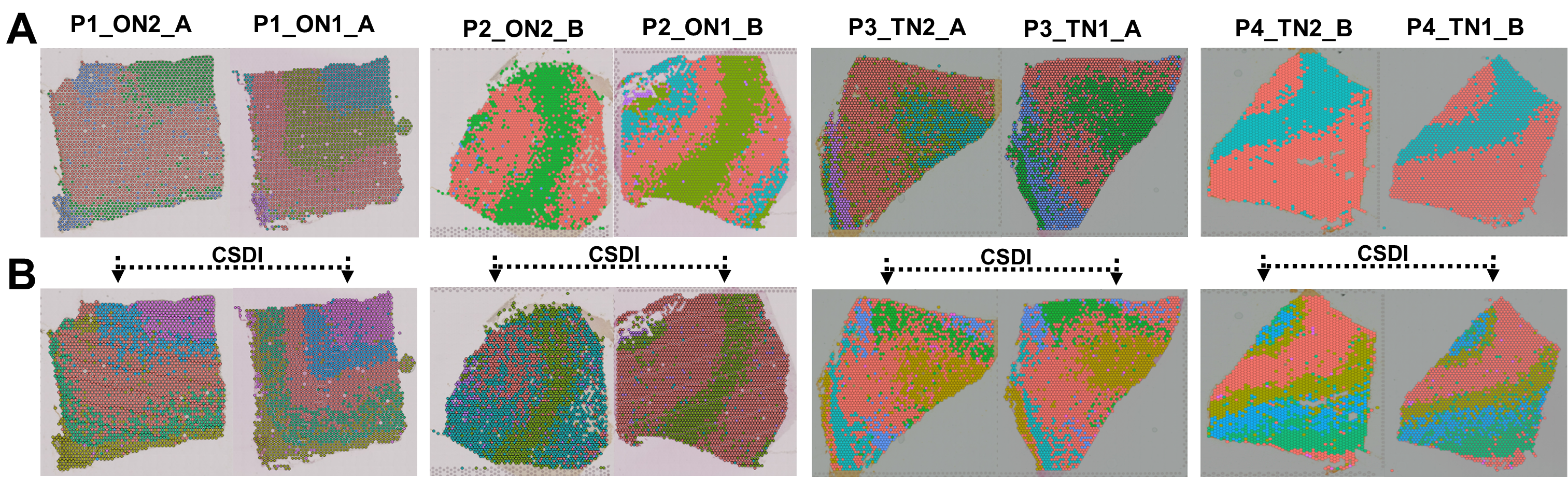

### Supplementary Figure 2

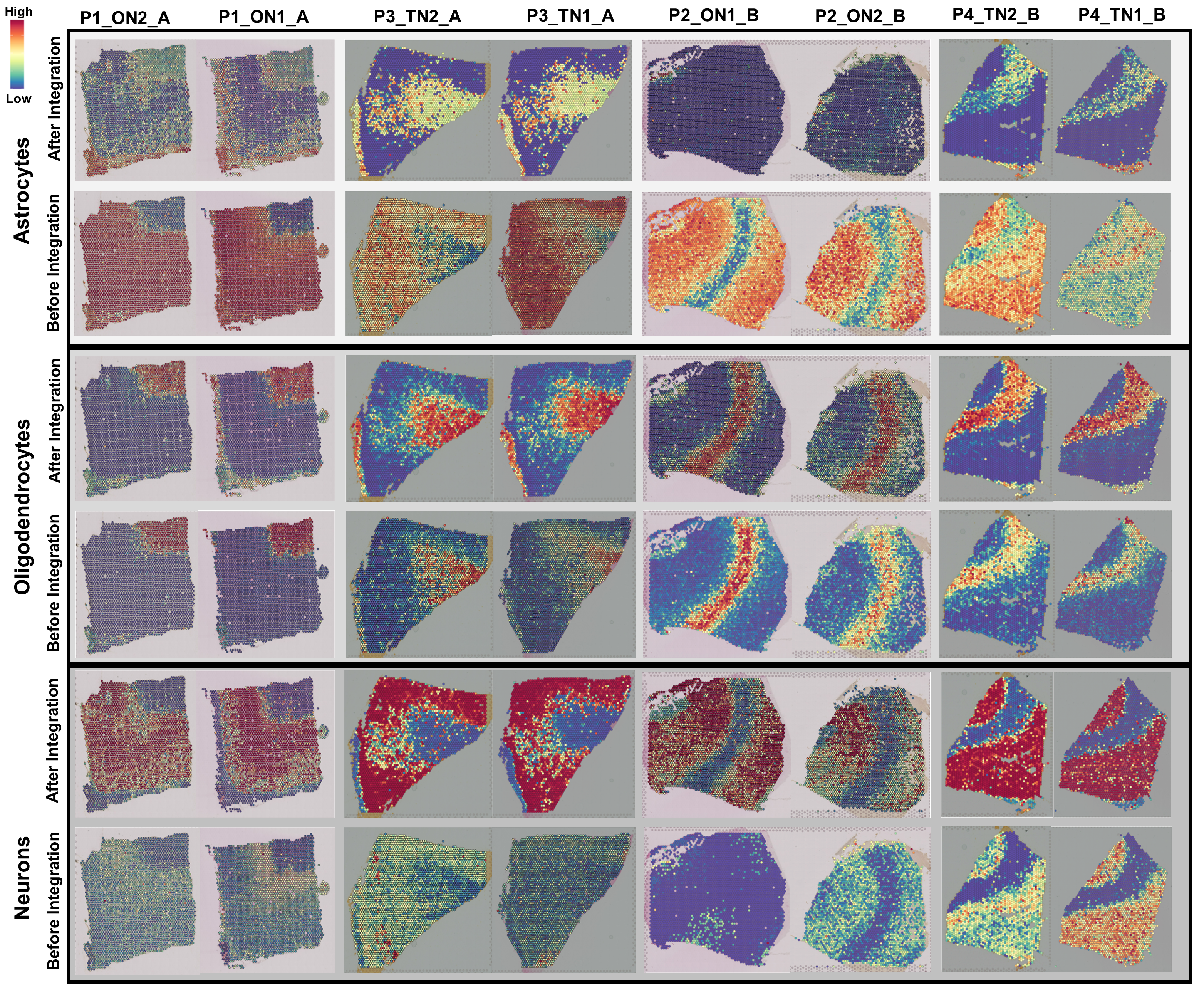

### Supplementary Figure 3

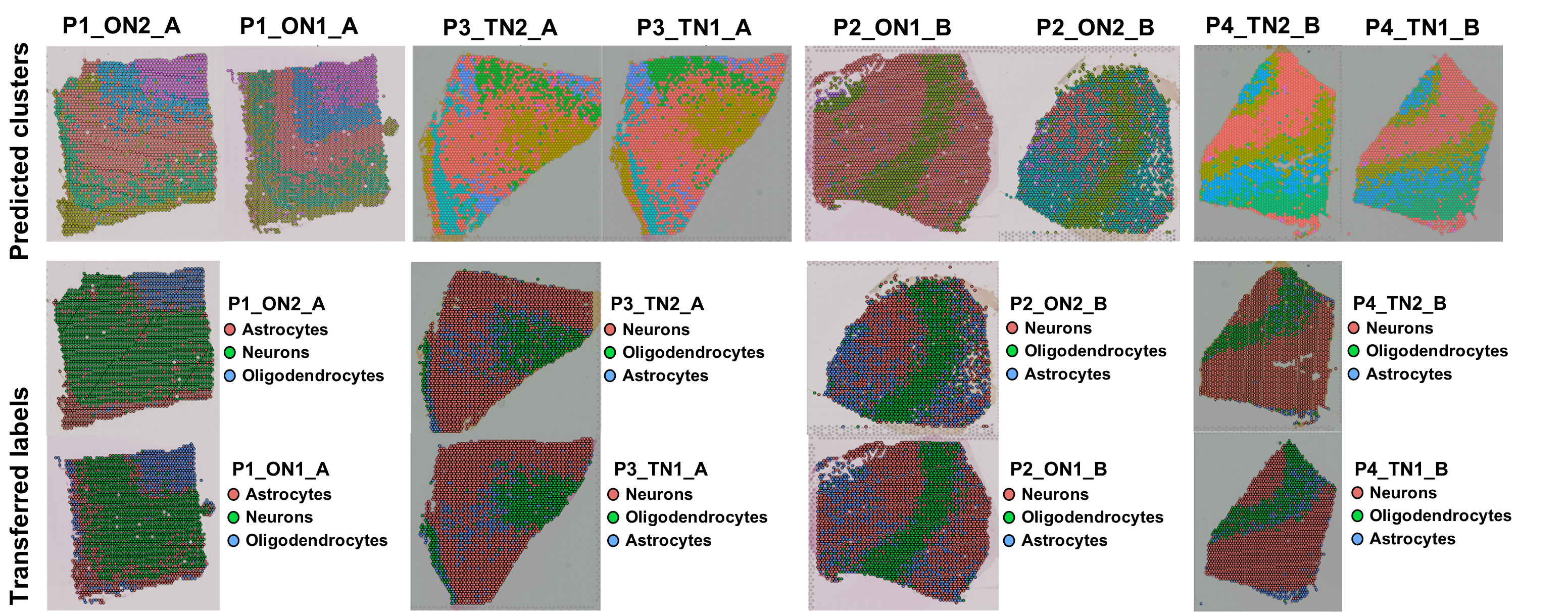
